# Hyaluronic Acid-Modified Mesoporous Manganese Dioxide Combined with Carboplatin Synergistically Enhances Ovarian Cancer Therapy

**DOI:** 10.1101/2025.08.07.669238

**Authors:** Yong Wang, Jie Pi, Yuzi Zhao, Qingzhen Xie

## Abstract

Ovarian cancer (OC) is a prevalent malignant tumor that poses a significant threat to women’s health. Conventional chemotherapeutic agents, such as carboplatin, are associated with several limitations, including notable side effects and the emergence of drug resistance. Carboplatin (CBP) and hyaluronic acid (HA) were subsequently loaded into nanoparticles through incubation to construct targeted nanoparticles (210.3 nm, -31.2 mV) capable of delivering both chemotherapy and chemodynamic therapy. In vitro experiments demonstrated that these nanoparticles enhanced ROS generation through enzyme-like activity and induced immunogenic cell death (ICD) by releasing damage-associated molecular patterns (DAMPs), such as ATP and HMGB1. In vivo animal studies revealed that the nanoparticles predominantly accumulated at the tumor site within 24 hours and significantly suppressed tumor growth by activating the cGAS-STING pathway. These findings indicate that the CBP@HMnO_2_@HA nanoparticles exhibit potent chemotherapeutic and chemodynamic properties, along with strong catalytic activity. This study presents a promising nanotechnology-based strategy for cancer therapy.

**Highlights:**

1. The CBP@HMnO_2_@HA nanoparticles integrate chemotherapy with chemodynamic therapy, thereby enhancing anti-tumor efficacy.
2. These nanoparticles induce ROS-mediated immunogenic cell death, leading to the release of DAMPs, including ATP and HMGB1.
3. Through HA-mediated targeting, the nanoparticles demonstrate preferential accumulation within tumor tissues, resulting in significant tumor growth inhibition in vivo while reducing off-target toxicity.

## 1. Introduction

Ovarian cancer (OC) poses a significant threat to women’s health globally. According to global cancer statistics, OC ranks first among gynecological malignancies in terms of both incidence and mortality. Its high recurrence rate and low survival rate present major challenges in clinical management [1, 2]. Conventional treatment strategies for ovarian cancer primarily involve surgical resection, chemotherapy, and radiotherapy; however, these approaches are often associated with considerable side effects and the development of drug resistance, which limit their long-term efficacy [3, 4]. Therefore, it is crucial to explore novel therapeutic strategies that are highly effective and minimally toxic.

Carboplatin (CBP), a second-generation platinum-based chemotherapeutic agent, exhibits reduced nephrotoxicity and fewer gastrointestinal side effects compared to cisplatin. It shares a similar mechanism of action with cisplatin but demonstrates superior antitumor activity [5]. CBP is predominantly used in the palliative treatment of advanced epithelial ovarian cancer, fallopian tube cancer, and primary peritoneal cancer [6-8]. Additionally, it is employed in the treatment of small cell lung cancer and head and neck squamous cell carcinoma [9, 10]. Previous studies have demonstrated that carboplatin induces DNA damage, thereby activating both the classical STING/TBK1/IRF3 pathway and the non-classical STING-NF-κB signaling complex [11]. This DNA damage also triggers an intracellular stress response, leading to immunogenic cell death (ICD). During ICD, cells release damage-associated molecular patterns (DAMPs), such as calreticulin (CRT), high mobility group box 1 (HMGB1), and adenosine triphosphate (ATP). The DNA damage caused by carboplatin enhances DAMP release, activates dendritic cells (DCs), promotes antigen presentation, and stimulates immune activation [12, 13].

Hollow mesoporous MnO_2_ (HMnO_2_), a nanomaterial with unique physicochemical properties, has shown promising applications in drug delivery, imaging diagnostics, and cancer therapy due to its high specific surface area, excellent biocompatibility, and catalytic performance [14-17]. Recent studies have demonstrated that Mn^2+^ are capable of inducing ICD through the release of DAMPs [18, 19]. These DAMPs, released during cellular stress or damage, are captured by antigen-presenting cells, thereby enhancing T-cell priming and activating anti-tumor immunity. Notably, manganese dioxide can be reduced to divalent Mn^2+^ in the tumor microenvironment, where excessive glutathione (GSH) is present [20]. Under the NaHCO3 buffering system, Mn^2+^ mediates the Fenton-like reaction with H_2_O_2_, effectively generating hydroxyl radicals (•OH), which are cytotoxic free radicals capable of inducing ICD and suppressing tumor growth [21, 22].

Moreover, Mn^2+^ not only directly activates the cyclic GMP-AMP synthase (cGAS)-stimulator of interferon genes (STING) pathway but also enhances STING activation via cGAMP [23]. Activation of the cGAS-STING pathway leads to the production of type I interferons (IFNs), thereby improving antigen presentation by DCs and augmenting tumor-specific T-cell responses [24]. Furthermore, HMnO_2_can catalyze the decomposition of excess H_2_O_2_ in the tumor microenvironment to generate oxygen, alleviating hypoxia and enhancing the efficacy of immunotherapy [25].

Based on these mechanisms, this study developed a tumor-targeted manganese-based nanoparticle formulation combined with chemotherapy and chemodynamic therapy to enhance the elimination of ovarian cancer cells through the induction of ICD and activation of the cGAS-STING pathway, as illustrated in Scheme 1. Mesoporous MnO_2_ was synthesized using silica nanoparticles as templates through a classical growth and etching process. To improve biocompatibility, targeting capability, and anti-tumor efficacy, carboplatin-loaded nanoparticles were further coated with hyaluronic acid (HA), forming a tumor-targeted nanomedicine (CBP@HMnO_2_@HA). HA not only enhances the biocompatibility of MnO_2_ nanoparticles but also specifically binds to receptors overexpressed on tumor cells, enabling targeted delivery. Upon internalization, HA is degraded, releasing carboplatin. Simultaneously, MnO_2_ is reduced to Mn^2+^, which catalyzes the Fenton-like reaction with H_2_O_2_ to produce •OH, significantly increasing ROS levels. This ROS elevation promotes ICD and facilitates tumor cell death. In addition, Mn^2+^ synergizes with carboplatin to activate the STING pathway, triggering potent anti-tumor immune responses. The innovation of this approach lies in the integration of chemotherapy and chemodynamic therapy, leveraging distinct yet complementary mechanisms to achieve enhanced tumor suppression. Carboplatin and Mn^2+^ work cooperatively to activate the STING pathway, thereby promoting tumor immunotherapy.

## 2. Materials and methods

### 2.1 Materials and reagents

Hyaluronic acid,potassium permanganate, sodium carbonate, tetraethylorthosilicate, carboplatin, methylene blue were purchased from Aladdin (Shanghai, China). Hydrogen peroxide was purchased from Sinopharm Group Chemical Reagent Co., LTD (Shanghai, China). All chemicals were obtained from commercial sources and used without further purification unless otherwise noted. DMEM medium with 4.5 g glucose, penicillin/streptomycin (P/S), 0.25% trypsin-EDTA, and Fetal Bovine Serum (FBS) were purchased from Gibco (Gran Island, NY, U.S.A.).

### 2.2 Cell lines and animals

All cells were purchased from the American Type Culture Collection (ATCC, Manassas, VA, USA).

BALB/c-nu (female, 7 weeks old) was purchased from Hunan SJA Laboratory Animal Co. Ltd. (Hunan, China) and raised in SPF animal rooms. All animal experiments were conducted in accordance with the guidelines of Hubei Provincial Center for Disease Control and Prevention Institutional Animal Care and Use Committee (202510069).

### 2.3 Preparation of CBP@HMnO_2_@HA

Silica nanoparticles (SiO_2_ NPs) were synthesized via an improved Stober method [26]. Specifically, 20 mL of deionized water, 10 mL of NH_3_·H_2_O, and 120 mL of ethanol were mixed in a 250 mL round-bottomed flask. The mixture was heated at 50°C in an oil bath under magnetic stirring for 15 minutes. Subsequently, 500 μL of tetraethylorthosilicate (TEOS) was added dropwise. The reaction mixture was stirred at 40°C for 8 hours to produce SiO_2_ nanoparticles. The resulting nanoparticles were sequentially washed with ethanol and water, then stored in triple-distilled water for further use. Under ultrasonic treatment (37°C, 40 kHz), 600 mg of potassium permanganate (KmnO_4_, 30 mg/mL) was slowly added dropwise to an aqueous solution, which was subsequently introduced into a suspension containing 50 mg of SiO_2_ nanoparticles. After overnight stirring at room temperature, mesoporous manganese dioxide-coated silica nanoparticles (SiO_2_@MnO_2_) were obtained by centrifugation at 12,000 rpm for 10 min. Hollow mesoporous manganese dioxide nanoparticles (HMnO_2_) were prepared by etching SiO_2_ from SiO_2_@MnO_2_. Specifically, the as-prepared mesoporous MnO2@SiO_2_was dispersed in a 2 M sodium carbonate (Na_2_CO_3_) aqueous solution and continuously stirred at 60°C for 12 h. The resultant mixture was centrifuged and washed multiple times with water to obtain HMnO_2_ nanoparticles. To fabricate CBP@HMnO_2_ nanoparticles, 20 mg of HMnO_2_ was added to 10 mL of phosphate-buffered saline (PBS) containing 4 mg of carboplatin (CBP), followed by 12 h stirring in the dark. The unbound CBP was removed by centrifugation, yielding CBP@HMnO_2_ nanoparticles. Finally, 10 mg of hyaluronic acid (HA) was added to a dispersion of CBP@HMnO_2_ nanoparticles in 10 mL of Tris-HCl buffer, and the mixture was stirred at room temperature in the dark for 6 h. After centrifugation, HA-functionalized CBP@HMnO_2_ nanoparticles (CBP@HMnO_2_@HA) were successfully obtained.

### 2.4 Characterization of CBP@HMnO_2_@HA

The morphology of the CBP@HMnO_2_@HA was firstly observed by transmission electron microscopy (TEM, HT7800, Hitachi, Japan). Dynamic Light Scattering (DLS, ZS90, Malvern, UK) was employed to measure their hydrodynamic diameter and potential. Next, the UV–vis absorbance data was collected using a UV–visible spectrometer (UV2600, Shimadzu, Japan). The encapsulation efficiency (EE%) and drug loading (DL%) of CBP were quantitatively analyzed using UV spectroscopy. The formulas for calculating EE% and DL% are as follows:

EE% = (Mass of encapsulated drug / Total drug mass) × 100%

DL% = (Mass of encapsulated drug / (Total mass of encapsulated drug + Carrier mass)) × 100%

### 2.5 Fenton-like reaction activity of CBP@HMnO_2_@HA

Methylene blue (MB) serves as a typical indicator for detecting •OH. Due to the strong oxidizing property of •OH, MB can be degraded, and the concentration of •OH is determined by monitoring the changes in UV spectral absorption of MB. In the experiment, 25 mM NaHCO_3_ buffer solution containing HA@HMnO_2_@CBP ([Mn^2+^] = 0.5 mM) and varying concentrations of GSH (0, 1, 2, and 5 mM) was incubated in a constant-temperature shaker at 37 °C for 15 min. Subsequently, 10 μg/mL MB solution and 10 mM H_2_O_2_ were added, and the mixture was further incubated for 30 min. The changes in the absorption peak of MB at 665 nm were measured using UV full-wavelength scanning.

### 2.6 The generation of dissolved oxygen in CBP@HMnO_2_@HA nanoparticles

An accurate preparation of 20 mL PBS buffer solution (pH 6.5) containing 500 μM H_2_O_2_ was performed, and four groups of this solution were prepared. Appropriate volumes of the final nanomedicine system solution (0, 25, 50, and 100 μg/mL) were sequentially added to each group of anoxic H_2_O_2_ solutions, followed by the rapid addition of 1 mL of HA@HMnO_2_@CBP solution at each respective concentration. The solutions were sealed with liquid paraffin and stirred for 5 min. During this period, a dissolved oxygen meter (JPSJ-605F, Leici, Shanghai) was used to measure and record the dissolved oxygen levels in real time at 30 s intervals. Additionally, the concentration of fixed nanoparticles was maintained at 50 μg/mL, and oxygen generation was detected under different hydrogen peroxide concentrations (0.5, 1, and 5 mM). Finally, an oxygen generation curve was constructed, with the abscissa representing the oxygen production time and the ordinate representing the amount of oxygen produced.

### 2.7 In vitro cellular experiments

#### 2.7.1 In vitro cellular uptake behavior

To investigate the cellular uptake behavior, ID-8 cells (5 × 10^4^ cells per well) were placed in 12-well plates and cultured for 24 h. Subsequently, Cy5.5@HMnO_2_@HA was added and incubated for different times (4, 8, 12 h). Then the cells were washed with PBS and gathered for confocal laser scanning microscope (CLSM, FV3000, Olympus Corporation, Japan). The Cell nucleus is strained by DAPI with blue fluorescence. The red fluorescence comes from Cy5.5.

#### 2.7.2 In vitro cytotoxicity assays

ID-8 cells (5 × 10^3^ cells per well) were placed in 96-well plates and cultured for 24 h. Subsequently, different concentrations of CBP (Carboplatin), HMnO_2_, CBP@HMnO_2_ and CBP@HMnO_2_@HA were added and incubated. After 24 h, replace the drug-containing medium with fresh medium supplemented with 10.0 μL of Cell Counting Kit-8 (CCK-8), and incubate the samples for an additional 2 h. Upon completion of the incubation period, transfer the 96-well plate to the microplate reader (Multiskan FC, Thermo Fisher Scientific, USA) and determine the OD value at a wavelength of 450 nm.

#### 2.7.3 In vitro apoptosis assays

The flow cytometry technique was employed to assess the apoptosis rate of cells. ID-8 cells (5 × 10^4^ cells per well) were plated in 12-well plates and cultured for 24 h. Subsequently, they were stimulated with CBP, HMnO_2_, CBP@HMnO_2_, and CBP@HMnO_2_@HA at a concentration of 2 μg/mL for CBP. After an incubation period of 36 h, the cells were collected, washed with PBS, and stained with Annexin V-FITC and PI for flow cytometric analysis.

#### 2.7.4 In vitro live/dead staining assay

ID-8 cells (5 × 10^4^ cells per well) were seeded in 12-well plates and cultured for 24 h. Subsequently, the cells were stimulated with CBP, HMnO_2_, CBP@HMnO_2_, and CBP@HMnO_2_@HA. After a 12 h incubation period, live cells (stained green) and dead cells (stained red) were distinguished using Calcein-AM (2 μmol/L) and propidium iodide (PI, 4 μmol/L), respectively. The stained cells were then visualized under an inverted fluorescence microscope (CLSM, IX73, Olympus Corporation, Japan).

#### 2.7.5 In vitro ROS generation assay

The cultured ID-8 cells were digested by trypsin, collected by centrifugation, and resuspended in 1 mL of culture medium. Subsequently, the cell suspension was rapidly counted, and an appropriate volume was seeded into confocal culture dishes at a density of 2×10^5^ cells per dish. After incubation for 24 h in a sterile incubator at 37 °C and 5% CO_2_, 10 μL of 5 μg/mL CBP, HMnO_2_, CBP@HMnO_2_, and CBP@HMnO_2_@HA solution was added to the cultures and incubated with the cells for 4 h. Following this, the cells were stained with DCFH-DA and Hoechst 33342 for 30 min, washed 3 times with PBS, and subsequently imaged using an inverted fluorescence microscope (CLSM, IX73, Olympus Corporation, Japan).

#### 2.7.6 In vitro ICD response assay

ID-8 cells (5 × 10^4^ cells per well) were seeded in a 6-well plate and cultured for 24 h. Subsequently, CBP, HMnO_2_, CBP@HMnO_2_, and CBP@HMnO_2_@HA was added to the wells, and the cells were further cultured for an additional 24 h. The culture medium was aspirated, and 200 μL of lysis buffer was added to each well. To ensure complete lysis, the lysis buffer was mixed thoroughly by pipetting or gently shaking the plate, ensuring full contact between the buffer and the cells. Typically, cell lysis occurs immediately upon exposure to the lysis buffer. Following lysis, the samples were centrifuged at 12,000 g for 5 min at 4 °C, and the supernatant was collected for subsequent determination of ATP, HMGB1, IFN-β, IL-12 and TNF-α levels (Flexstation3, Molecular Devices, USA). Thereafter, the cells were washed with PBS and incubated with Alexa Fluor 647-conjugated anti-mouse CRT antibody in the dark for 45 min. The antibody solution was aspirated, and the cells were washed twice with PBS. Trypsin digestion was performed for 3 min, followed by stopping the digestion process. The cells were then resuspended, centrifuged at 450 g for 5 min, and washed twice with PBS to obtain the final cell pellet. The expression of CRT on the cell surface was analyzed using flow cytometry (CytoFLEX, BECKMAN, USA).

### 2.7 In vivo assessment

#### 2.7.1 In vivo biodistribution assay

HMnO_2_ and HMnO_2_@HA labeled with Cy5.5 (Cy5.5@HMnO_2_, Cy5.5@HMnO_2_@HA) was used to investigate the biodistribution in vivo. Female BALB/c-nu mice were subcutaneously injected with ID-8 cells (1 × 10^6^ cells per mouse) in the left axilla to construct ovarian-tumor mice models. After two weeks, tumor mice were randomly divided into two groups (n = 3), and they were injected via tail vein with Cy5.5@HMnO_2_ and Cy5.5@HMnO_2_@HA (Cy5.5 =0.5 mg/kg), respectively. The in vivo fluorescence images were obtained at 4, 6, 8, 12 and 24 h post injection by using a fluorescence living imaging approach (LIVIS Lumina III, PerkinElmer, USA). After 24 h post-injection, tumors and major organs were excised for ex vivo fluorescence imaging.

#### 2.7.2 In vivo antitumor efficacy

When the tumor reached a volume of 50 mm^3^, the mice were randomly assigned into five groups (Group 1: Saline, Group 2: CBP, Group 3: HMnO_2_, Group 4: CBP@HMnO_2_, Group 5: CBP@HMnO_2_@HA), with three mice in each group [27]. The mice were administered treatment via tail vein injection every two days for a total of five doses. The dosage was determined as 3.5 mg/kg based on the CBP concentration. The initial administration was designated as Day 0. Body weight and tumor volume were measured every other day throughout a continuous 14-day observation period. At the conclusion of the 14-day observation, the mice were euthanized, and their organs and tissues as well as tumors were excised. The weights of these tissues and tumors were recorded, and images of the tumors were captured. The volume was calculated using the following formula: V = a (Long diameter) *b (Short diameter) ^2^/2. Mice in each group were killed on the 14th day. Blood samples were gathered for blood routine and biochemistry analysis. Part of the tumors and major organs were harvested for hematoxylin and eosin (H&E), Ki67, and terminal deoxynucleotidyl transferase-mediated deoxyUTP-biotin nick end labeling (TUNEL) staining.

#### 2.7.3 Western blotting

Tumors harvested from mice post-treatment were lysed with RIPA lysis buffer (Beyotime) to extract total protein. Protein concentrations were quantified using the BCA Protein Assay Kit (Beyotime). Equal amounts of protein from each group were resolved by SDS-PAGE and subsequently electrotransferred onto PVDF membranes. The membranes were then blocked with 5% non-fat milk and incubated overnight at 4°C with specific primary antibodies, such as anti-GAPDH (A19056, ABclonal), anti-p-STING (A21051, ABclonal), anti-TBK-1 (A3458, ABclonal), anti-p-TBK-1 (AP1026, ABclonal), anti-IRF-3 (A11118, ABclonal), anti-p-IRF-3 (AP0995, ABclonal). Following incubation with the secondary antibody, the proteins were visualized using ECL Substrate. Semi-quantitative analyses were performed using Image J software.

#### 2.7.4 In vivo antitumor immune response

The immune-related cytokines from tumors including IFN-β, TNF-α, and IL-12 were determined by ELISA kits, and the specific operation methods were carried out according to standard protocols.

### 2.8 Statistical analysis

Statistical analysis was performed using one-way analysis of variance (ANOVA) or t-test for multiple comparisons. All data were presented as mean ± standard deviation (SD). Statistical significance was set at the following levels: *p < 0.05, **p < 0.01, ***p < 0.001, and ****p <0.0001.

## 3. Results

### 3.1 Preparation and characterization of NPs

Firstly, silica nanoparticles were synthesized using the Stober method, and HMnO_2_ nanoparticles were prepared via a reduction reaction followed by potassium permanganate etching. Subsequently, CBP and HA were loaded onto the nanoparticles through an incubation approach to fabricate targeted nanoparticles, CBP@HMnO_2_@HA, for combined chemotherapy and chemodynamic therapy, as illustrated in Fig. 1A. As shown in Fig. S1, DLS analysis revealed that the average hydrodynamic diameter of SiO_2_ nanoparticles was 122.1 nm with a zeta potential of -31.5 mV, while that of SiO_2_@MnO_2_ was 134.0 nm with a zeta potential of -30.3 mV. TEM images confirmed that all nanoparticles exhibited a spherical morphology, indicating successful synthesis. Notably, a distinct HA surface layer was observed in CBP@HMnO_2_@HA, confirming effective HA coating (Fig. 1A). Further characterization by DLS showed that the mean particle sizes of HMnO_2_, CBP@HMnO_2_, and CBP@HMnO_2_@HA were 168.0 nm, 191.0 nm, and 210.3 nm, respectively (Fig. 1B). While no significant size difference was observed between HMnO_2_ and CBP@HMnO_2_, a notable increase was found between HMnO_2_ and CBP@HMnO_2_@HA, likely due to HA coating. The mean zeta potentials of HMnO_2_ (−30.0 mV), CBP@HMnO_2_ (−30.5 mV), and CBP@HMnO_2_@HA (−31.2 mV) remained relatively consistent; however, the slightly increased negative charge in CBP@HMnO_2_@HA may be attributed to HA modification. Importantly, all samples exhibited absolute zeta potentials exceeding 30 mV (Fig. 1C), suggesting good colloidal stability, and PDI values were below 0.3, indicating uniform dispersion (Fig. S2). Based on the carboplatin standard curve (Fig. S3), the drug loading content of CBP in CBP@HMnO_2_@HA and CBP@HMnO_2_ were determined to be 6.32% and 9.26%, respectively, with corresponding encapsulation efficiency detailed in Table S1.

**Figure 1.**
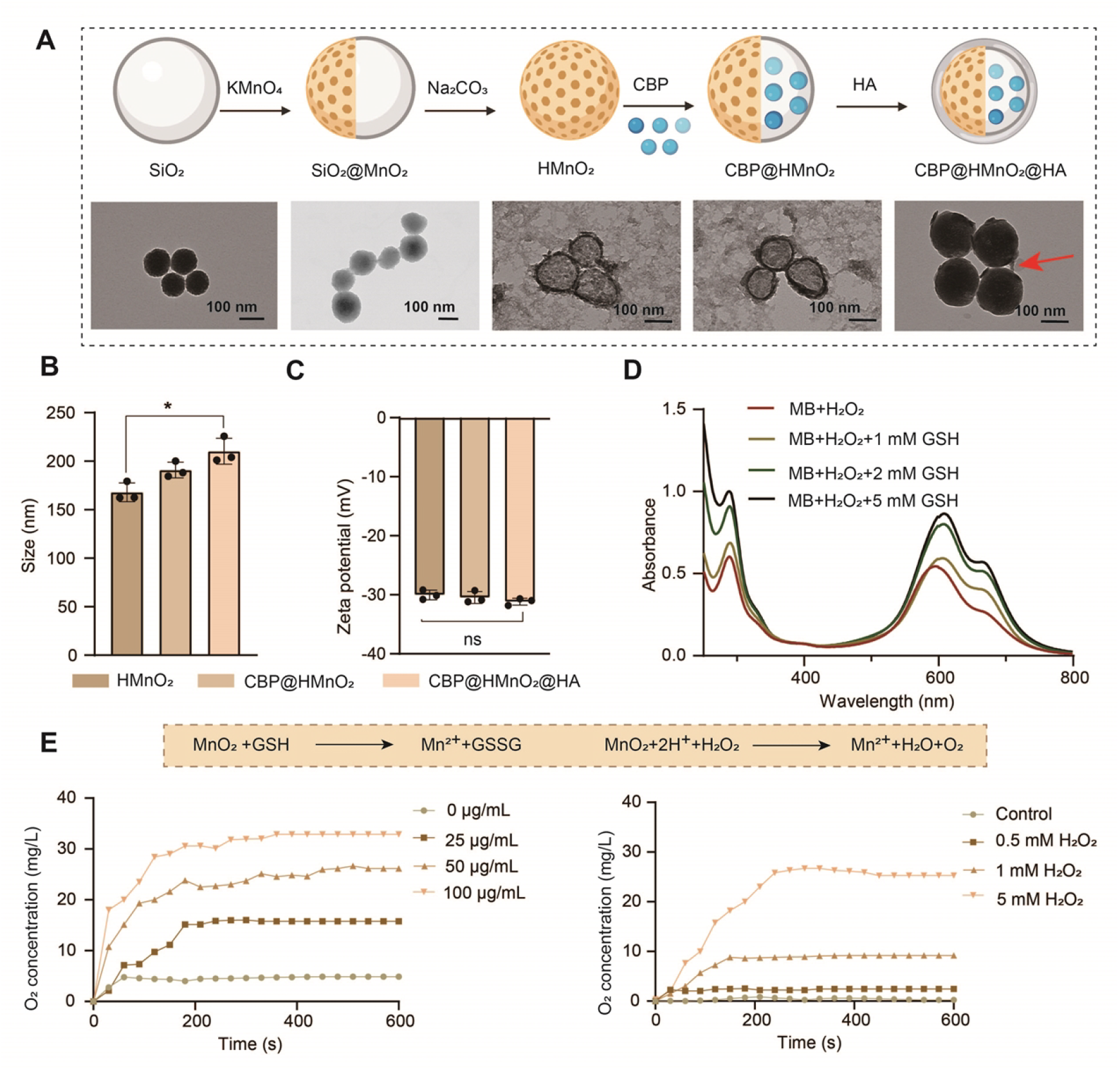
Preparation and characterization of NPs. (A) The flowchart of the preparation of nanoparticles and representative TEM image of NPs (scale bars represented 100 nm). (B) Hydrodynamic diameters of NPs by DLS. (C) Potential of NPs by DLS. (D) The Fenton-like reaction ability from NPs in the presence of 1 mM, 2 mM, 5 mM GSH and H_2_O_2_ at 37^°^C, respectively. (E) Characterization diagram of the ability of nanoparticles of different concentrations to catalyze the generation of O_2_ from different concentrations of H_2_O_2_. Data are presented as mean ± SD, n=3. Statistical significances between every two groups were calculated via one-way ANOVA. *p < 0.05.

Methylene blue (MB) was employed to evaluate the chemodynamic performance of manganese dioxide [28].

These results demonstrate that Mn^2+^ within the carrier exhibit efficient glutathione (GSH)-depletion capability, reacting with GSH to generate Mn^2+^, which subsequently facilitates a Fenton-like reaction with H_2_O_2_ to produce hydroxyl radicals. To assess the oxygen-generating capacity of CBP@HMnO_2_@HA, varying concentrations of CBP@HMnO_2_@HA were introduced into H_2_O_2_solutions. At 100 μg/mL of CBP@HMnO_2_@HA, dissolved oxygen levels reached 32.8 μg/mL after 600 s (Fig. 1E). Similarly, when different H_2_O_2_ concentrations were applied to CBP@HMnO_2_@HA, 5 mM H_2_O_2_yielded 25.3 μg/mL dissolved oxygen at 600 s (Fig. 1E). Compared to the H_2_O_2_control group, CBP@HMnO_2_@HA group significantly accelerated oxygen generation, with dissolved oxygen concentration increasing steadily over time. These findings indicate that CBP@HMnO_2_@HA group serves as an effective oxygen-delivery agent, capable of alleviating tumor hypoxia and supplying sufficient oxygen for chemodynamic therapy, thereby facilitating a cascade catalytic process.

### 3.2 In vitro anticancer effects of NPs

To evaluate the in vitro anti-tumor efficacy of the nanoparticles, we conducted a series of experiments on ID-8 cells (Fig. 2A). The cells were treated with varying concentrations (0–20 μg/mL) of CBP, HMnO_2_, CBP@HMnO_2_, and CBP@HMnO_2_@HA nanoparticles. The CCK-8 assay results demonstrated that at a concentration of 20 μg/mL, the cell viability rates were 64.0%, 59.5%, 31.1%, and 17.8% for CBP, HMnO_2_, CBP@HMnO_2_, and CBP@HMnO_2_@HA, respectively (Fig. 2B). Notably, the anti-tumor effects of the CBP and HMnO_2_ groups were comparable. CBP exerts its cytotoxic effect through chemotherapy, whereas HMnO_2_ induces tumor cell death via chemodynamic therapy by generating reactive oxygen species (ROS). Furthermore, the IC50 values were determined as 12.7 μg/mL for CBP@HMnO_2_ and 5.8 μg/mL for CBP@HMnO_2_@HA (Fig. 2C), indicating that CBP@HMnO_2_@HA exhibited the strongest anti-proliferative activity, which may be attributed to the tumor-targeting capability of HA.To visually assess the inhibitory effects of the nanoparticles on ID-8 cells, live/dead staining was performed (Fig. S4). The fluorescence imaging results were consistent with those obtained from the cytotoxicity and apoptosis assays, further supporting the superior anti-proliferative effect of CBP@HMnO_2_@HA.

**Figure 2.**
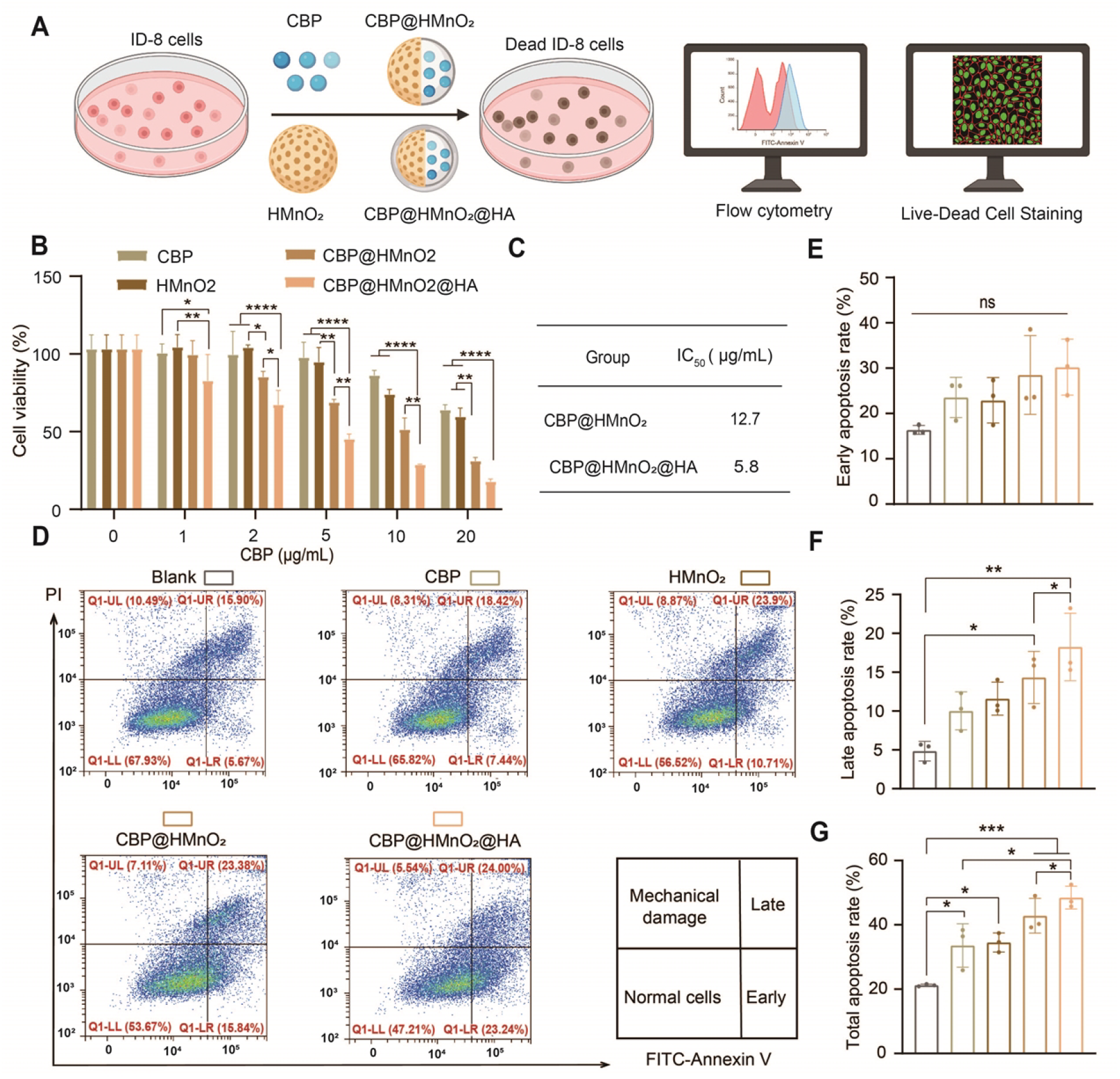
*In vitro* anticancer effects of NPs. (A) Schematic representation of the in vitro cell experiment procedure. (B) Relative cell viabilities of ID-8 cells treated with CBP (Carboplatin), HMnO_2_, CBP@HMnO_2_ and CBP@HMnO_2_@HA NPs via an CCK-8 assay at 24 h, respectively. (C) The IC50 values of various nanoparticles against ID8 cells. (D) Apoptosis rate and (E-G) auantification of apoptosis via FCM on ID-8 cells treated with CBP, HMnO_2_, CBP@HMnO_2_ and CBP@HMnO_2_@HA NPs at 24 h. Data are presented as mean ± SD, n=3. Statistical significances between every two groups were calculated via one-way ANOVA. *p < 0.05, **p < 0.01, ***p < 0.001.

To investigate the mechanism of nanoparticle-induced cell death, flow cytometry was employed to analyze apoptosis levels (Fig. 2D). Although no significant differences were observed in early apoptosis among the treatment groups compared to the Blank group (Fig. 2E), notable differences were found in late apoptosis. Specifically, the late apoptosis rates were 5.1% (Blank), 9.8% (HMnO_2_), 13.4% (CBP@HMnO_2_), and 18.1% (CBP@HMnO_2_@HA) (Fig. 2F). These findings suggest that the nanoparticles primarily induce tumor cell death through late apoptosis.In terms of total apoptosis rate (Fig. 2G), all treatment groups showed apoptotic effects, with CBP@HMnO_2_@HA demonstrating significantly higher efficacy compared to both the CBP and HMnO_2_ groups. This result highlights the synergistic effect of CBP and HA in enhancing therapeutic outcomes.

### 3.3 In vitro ICD effects of NPs

Fig. 3A presents a schematic illustration of the activation of the immunogenic cell death (ICD) effect through ROS generation by nanoparticles *in vitro*. To evaluate nanoparticle uptake, laser confocal imaging was performed at 4, 8, and 12 h. The results demonstrated that nanoparticle uptake increased progressively over time. The average fluorescence intensities for the Blank, Cy5.5@HMnO2, and Cy5.5@HMnO2@HA groups were 0.07, 2.18, and 2.55, respectively (Fig. 3B and Fig. S5). Both the Cy5.5@HMnO_2_ and Cy5.5@HMnO_2_@HA groups exhibited statistically significant differences compared to the Blank group. Moreover, a significant difference was observed between the Cy5.5@HMnO_2_ and Cy5.5@HMnO_2_@HA groups, indicating that HA coating enhances HMnO_2_ tumor targeting and improves cellular uptake efficiency. This finding also explains why CBP@HMnO_2_@HA exhibits greater anti-tumor efficacy than CBP@HMnO_2_. To assess the ROS-generating capacity of HMnO_2_, CLSM images were obtained (Fig. 3C). As shown in Fig. 3C, the average fluorescence intensities were as follows: CBP (0.08), HMnO2 (291.04), CBP@HMnO2 (334.90), and CBP@HMnO_2_@HA (2090.83). These results indicate that CBP@HMnO2@HA generates significantly more ROS, primarily due to enhanced cellular uptake.

**Figure 3.**
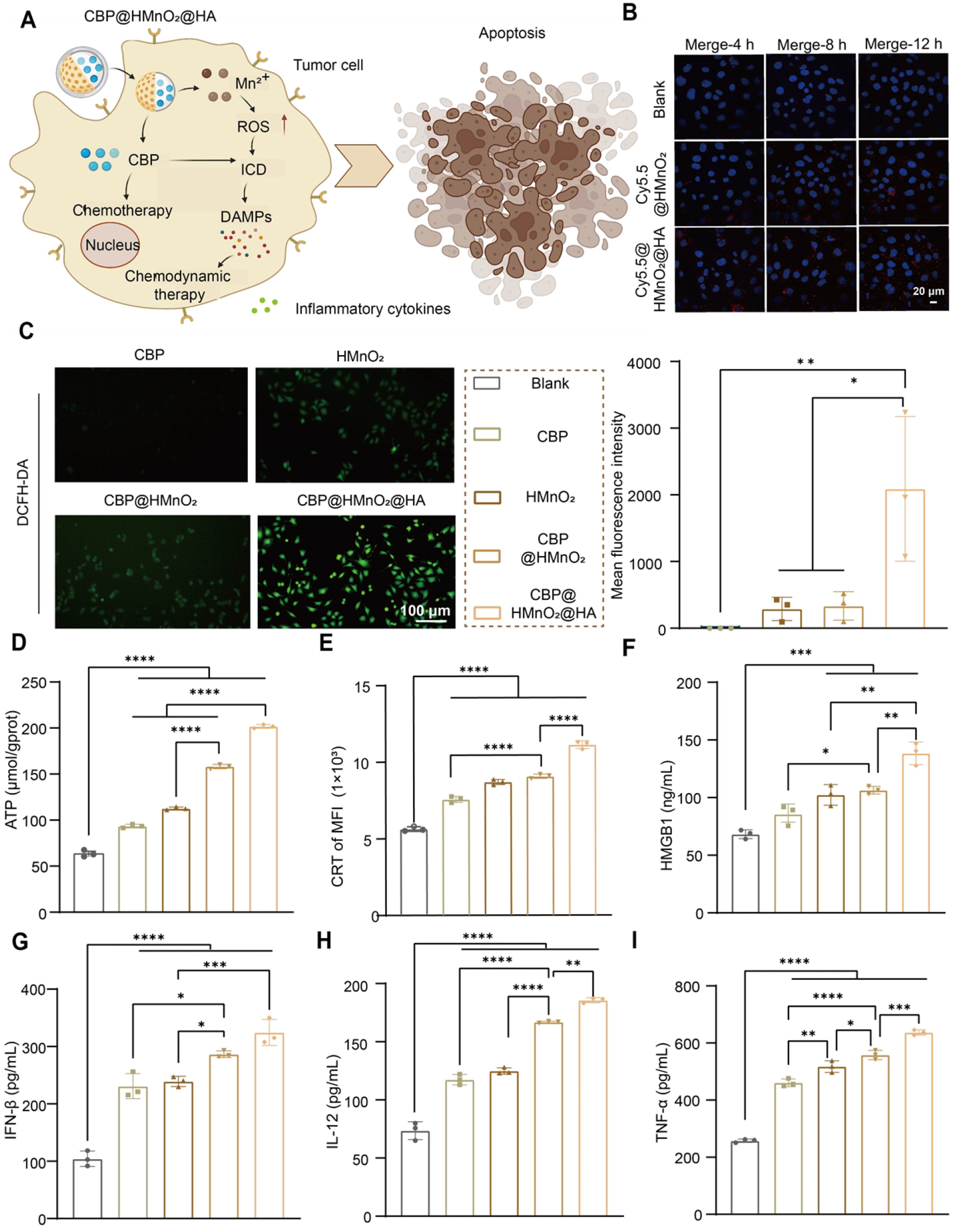
*In vitro* ICD effects of NPs. (A) Schematic diagram of the ICD effect activated by nanoparticles through the generation of ROS *in vitro*. (B) Representative CLSM images of ID-8 cells treated with Cy5.5@NPs at 4 h, 8 h and 12 h, respectively. The Cell nucleus is strained by DAPI with blue fluorescence. The red fluorescence comes from Cy5.5 (scale bar represented 20 μm). (C) Reactive oxygen species generation of ID-8 cells treated with CBP, HMnO_2_, CBP@HMnO_2_ and CBP@HMnO_2_@HA NPs at 24 h, respectively (Scale bar represented 100 μm). (D) The ATP content test kit is designed to measure the levels of ATP present within ID-8 cells. (E) CRT quantification of fluorescence intensity via FCM on ID-8 cells treated with CBP, HMnO_2_, CBP@HMnO_2_, and CBP@HMnO_2_@HA NPs. (F) The Elisa assay of ID-8 cells treated with various drugs to determinate HMGB1 content. The Elisa of ID-8 cells treated with various drugs to assay G) IFN-β, H) IL-12 and I) TNF-α. Data are presented as mean ± SD, n=3. Statistical significances between every two groups were calculated via one-way ANOVA. *p < 0.05, **p < 0.01, ***p < 0.001, ****p < 0.0001.

Within cells, manganese dioxide is reduced to Mn^2+^, which mediates a Fenton-like reaction to produce hydroxyl radicals (•OH), thereby effectively increasing intracellular ROS levels. This leads to the activation of the ICD pathway and subsequent tumor cell death. During ICD, cells release DAMPs, such as CRT, HMGB1, and ATP. Additionally, ICD promotes the secretion of pro-inflammatory cytokines. Therefore, we first measured ATP levels in the cell supernatant. As shown in Fig. 3D, the ATP content in the Blank, CBP, HMnO_2_, CBP@HMnO_2_, and CBP@HMnO_2_@HA groups was 63.24, 93.53, 112.70, 158.13, and 201.70 μmol/gprot, respectively. Flow cytometry was employed to quantify CRT expression in ID-8 cells treated with different nanoparticles. As illustrated in Fig. 3E, the average fluorescence intensities were as follows: Blank group (5641.67), CBP group (7570.03), HMnO_2_ group (8702.03), CBP@HMnO_2_ group (9069.43), and CBP@HMnO2@HA group (11125.60). The ELISA method was used to measure HMGB1 levels in ID-8 cells following treatment with various formulations (Fig. 3F-i). According to the data in Fig. 3F, the HMGB1 concentrations in the Blank, CBP, HMnO_2_, CBP@HMnO_2_, and CBP@HMnO_2_@HA groups were 68.30, 85.58, 102.33, 106.40, and 138.37 pg/mL, respectively.

Furthermore, Fig. 3G–I present the expression levels of pro-inflammatory cytokines (IFN-β, IL-12, and TNF-α) detected via Elisa. Overall, the results from Fig. 3D–I demonstrate that all treatment groups (CBP, HMnO_2_, CBP@HMnO_2_, and CBP@HMnO_2_@HA) showed statistically significant increases compared to the Blank group, confirming that both CBP and HMnO_2_ can induce the ICD effect. HMnO_2_ activates ICD-related markers through ROS production. Upon entering tumor cells, CBP forms Pt-DNA adducts, leading to tumor cell necrosis or apoptosis. During this process, tumor cells undergo stress responses, resulting in elevated surface expression of heat shock proteins (HSPs) and translocation of immune-related molecules such as calreticulin to the cell membrane. Simultaneously, cells release ATP, HMGB1, and immunostimulatory cytokines, further promoting ICD activation.

### 3.4 In vivo biodistribution of NPs

CD44 is a transmembrane glycoprotein that possesses a specific domain at the N-terminus of its extracellular functional region, enabling specific binding to HA [29]. Upon specific interaction between HA and the CD44 receptor, which is overexpressed on the surface of tumor cells, clathrin-mediated endocytosis is initiated [30]. As a result, HA and its associated nanocarriers are internalized and transported to lysosomes via endocytic pathways. This mechanism allows HA to precisely deliver therapeutic agents to tumor cells, thereby facilitating targeted tumor therapy. To assess the targeting capability of HA-modified nanoparticles for effective tumor suppression, we evaluated their tumor accumulation and biodistribution using *in vivo* fluorescence imaging in an ID-8 ovarian cancer tumor model. In vivo fluorescence signals of both formulations at various time points are presented in Fig. 4A-B. The results demonstrate that Cy5.5@HMnO_2_@HA exhibits significantly stronger fluorescence accumulation at the tumor site compared to Cy5.5@HMnO_2_. These findings suggest that the enhanced tumor localization of Cy5.5@HMnO_2_@HA is attributable to the HA-mediated targeting of CD44. *Ex vivo* analysis further confirmed higher fluorescence intensity specifically within the tumor tissue, with a more pronounced signal observed in the Cy5.5@HMnO_2_@HA group than in the Cy5.5@HMnO_2_ group (Fig. 4A and C). Moreover, *ex vivo* organ imaging revealed that both Cy5.5@HMnO_2_@HA and Cy5.5@HMnO_2_ were predominantly accumulated in major metabolic organs, particularly the liver. Collectively, these results indicate that Cy5.5@HMnO_2_@HA demonstrates promising tumor-targeting efficacy against ovarian cancer and holds potential for inhibiting the growth of solid tumors.

**Figure 4.**
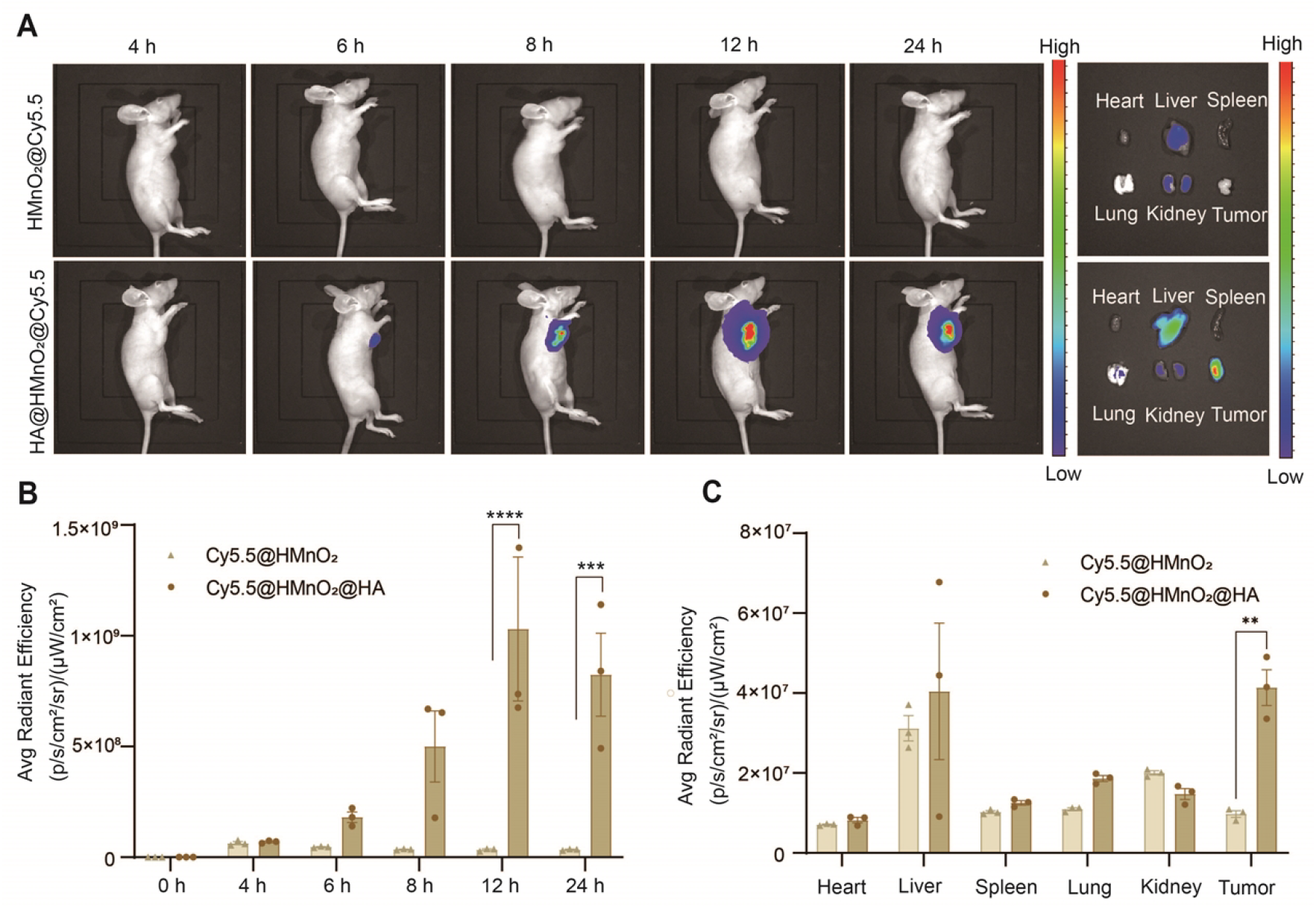
*In vivo* biodistribution of NPs. (A) *In vivo* fluorescent imaging of tumors at different times and *ex vivo* imaging of major organs and tumor at 24 h after intravenous injection of Cy5.5@NPs. (B) Quantification of average fluorescence intensity at tumor site at different time points and (C) the average fluorescence intensity of Cy5.5@NPs (Cy5.5@HMnO_2_ and Cy5.5@HMnO_2_ @HA) in major organs and tumor at 24 h after intravenous. Data are presented as mean ± SD, n=3. Statistical significances between two groups were calculated *via* t test. *p < 0.05, **p < 0.01, ***p < 0.001, ****p < 0.001.

### 3.5 In vivo antitumor effects of NPs

Given the potent antitumor activity observed in vitro and the tumor-targeting capability of CBP@HMnO_2_@HA, we aimed to evaluate its efficacy in inhibiting tumor growth using an ID-8 tumor-bearing mouse model. The specific treatment protocols are illustrated in Fig. 5A. Mice with tumors reaching approximately 50 mm^3^ were administered various treatments, including saline, CBP, HMnO_2_, CBP@HMnO_2_, and CBP@HMnO_2_@HA. As shown in Fig. 5B, no significant changes in body weight were observed during the treatment period. As depicted in Fig. 5C, the saline group exhibited the fastest tumor growth rate, whereas the free CBP group, HMnO_2_ group, and CBP@HMnO_2_ group all demonstrated significant tumor growth inhibition, which can be attributed to the established chemotherapeutic effect of CBP and the chemokinetic properties of HMnO_2_. Notably, CBP@HMnO_2_@HA exhibited enhanced anticancer efficacy, likely due to its prolonged circulation time in vivo and active tumor targeting. This supports the hypothesis that its antitumor effect results from the synergistic action of CBP, Mn^2+^, and HA. In the CBP@HMnO_2_@HA group, tumor growth was significantly slower, with a final average tumor volume of 145.43 mm^3^, compared to 823.80 mm^3^, 533.57 mm^3^, 454.83 mm^3^, and 357.47 mm^3^ in the saline, CBP, HMnO_2_, and CBP@HMnO_2_ groups, respectively. Tumor weights and images across the different treatment groups were consistent with the measured tumor volumes (Fig. 5D-E). Furthermore, immunofluorescence staining was conducted to assess Ki67, TUNEL expression and hematoxylin and eosin (H&E) staining of tumor tissues. The results indicated that CBP@HMnO_2_@HA exerted a marked inhibitory effect on tumor growth compared to the saline group, thereby providing additional evidence of its potent antitumor activity (Fig. 5F).

**Figure 5.**
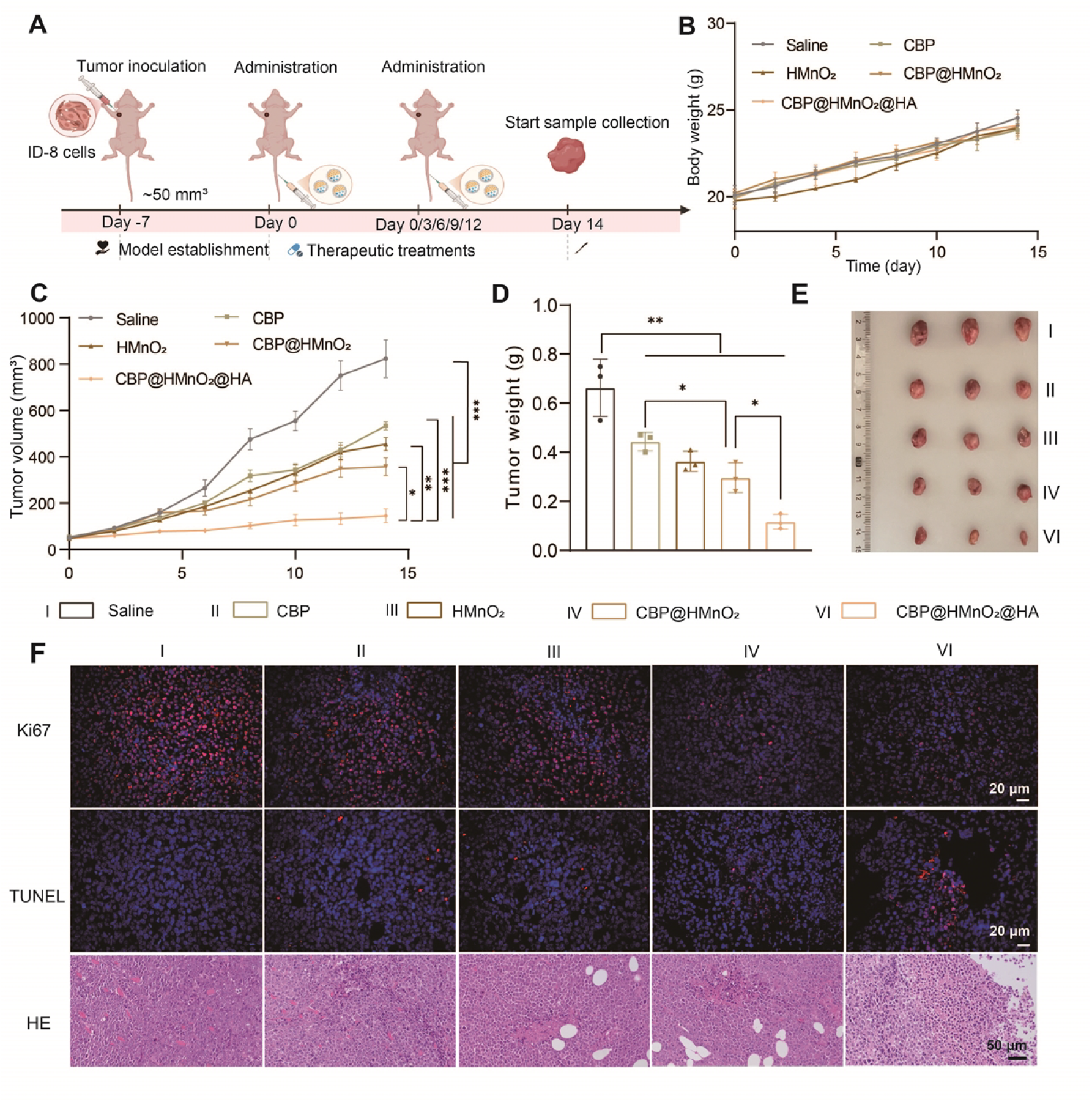
*In vivo* antitumor effects of NPs. (A) Schematic illustration of the treatment process. (B) Body weight changes of ID-8 tumor nude mice were recorded every two days for 14 days. (C) Relative tumor volumes of ID-8 tumor nude mice were recorded every two days. (D) Tumor weights and (E) corresponding tumor images of mice after different treatments. (F) TUNEL, Ki67 and H&E staining images of tumor slices excised from mice in different groups. Scale bars represented 20 μm, 20 μm and 50μm, respectively. Data are presented as mean ± SD, n=3. Statistical significances between every two groups were calculated *via* one-way ANOVA. *p < 0.05, **p < 0.01, ***p < 0.001.

### 3.6 In vivo activation of the cGAS-STING pathway

We previously reported that CBP can induce DNA damage and activate the classical STING/TBK1/IRF3 signaling pathway. Additionally, Mn^2+^ has also been shown to activate the cGAS-STING pathway. Activation of this pathway leads to the production of type I interferons (IFNs), which in turn enhances the expression of various proinflammatory cytokines, as illustrated in Fig. 6A. To further elucidate the roles of Mn^2^+ and CBP in the activation of the cGAS-STING pathway by CBP@HMnO2@HA, we performed Western blot analysis to evaluate the expression levels of p-STING, TBK1, p-TBK1, IRF3, and p-IRF3—key downstream effector molecules of the cGAS-STING pathway (Fig. 6B). As anticipated, free CBP and manganese ions alone induced only weak phosphorylation of STING and IRF3 proteins. In contrast, treatment with CBP@HMnO_2_@HA significantly enhanced the phosphorylation of these downstream signaling proteins, resulting in reduced tumor cell proliferation. This observation reflects the synergistic enhancement of the antitumor efficacy of CBP@HMnO_2_@HA (Fig. 6C– E). Furthermore, CBP@HMnO_2_@HA markedly promoted IFN-β production (Fig. 6F), confirming its ability to effectively activate the cGAS-STING pathway, as IFN-β secretion is a well-established marker of STING activation. Notably, CBP@HMnO_2_@HA also significantly increased the secretion of IL-12 and TNF-α, indicating that the nanoparticles can modulate the tumor microenvironment and exert potent antitumor effects. Collectively, these findings demonstrate that CBP@HMnO_2_@HA-based nanotherapy effectively activates the cGAS-STING pathway, thereby synergistically suppressing ovarian cancer progression.

**Figure 6.**
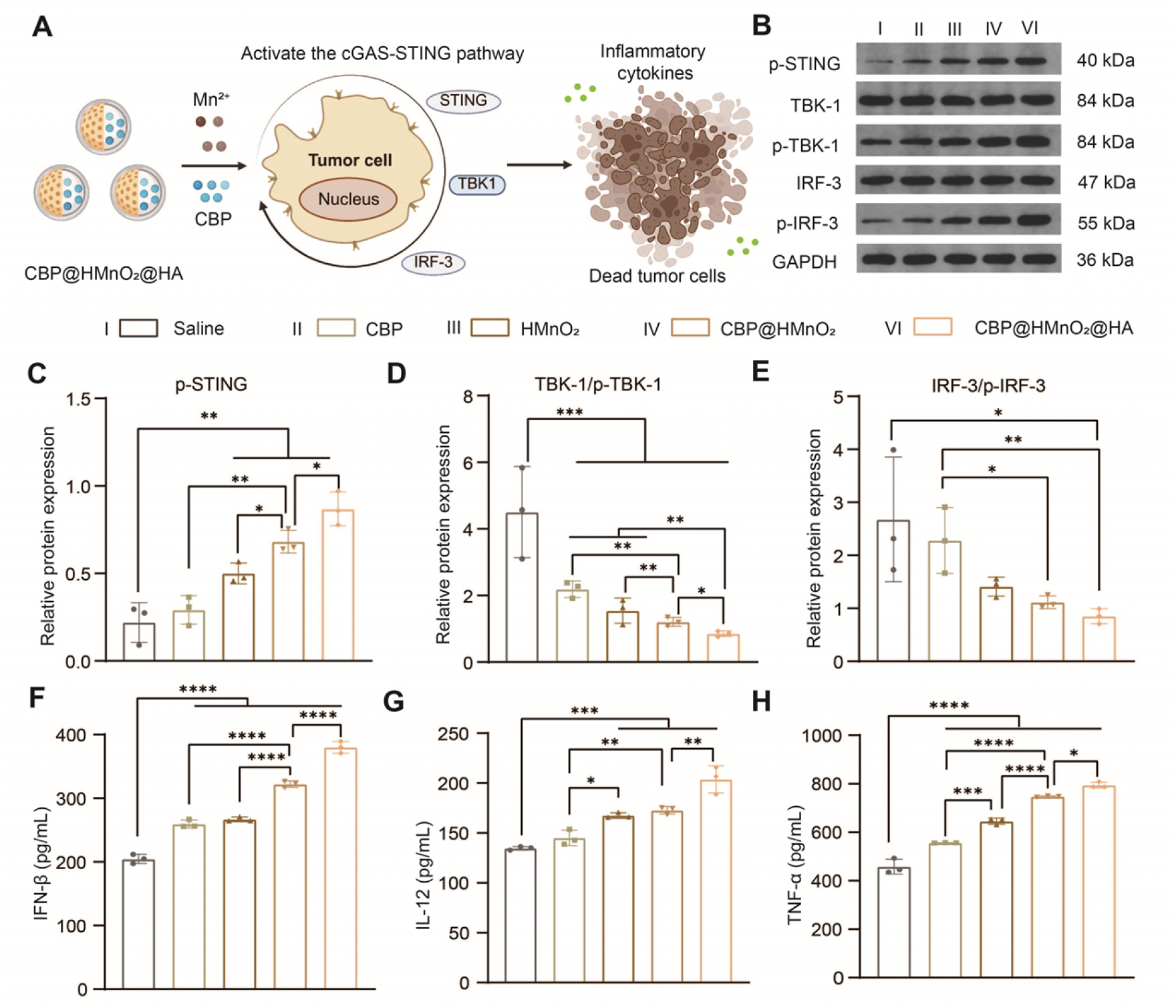
*In vivo* activation of the cGAS-STING pathway. (A) Schematic illustration of the activation process. (B) Western blot images and semi-quantitative analysis of cGAS-STING-related proteins (C-E) after different treatments. Cytokine levels of IFN-β (F), IL-12 (G), and TNF-α (H) in tumor tissues. Data are presented as mean ± SD, n=3. Statistical significances between every two groups were calculated *via* one-way ANOVA. *p < 0.05, **p < 0.01, ***p < 0.001, ****p < 0.0001.

### 3.7 In vivo biosafety evaluation of NPs

To evaluate the safety of the nanoparticles, we conducted hemolysis assays across multiple organs. The results presented in Fig. S6 demonstrate that, in comparison with the positive control group, various concentrations of CBP and CBP@HMnO_2_@HA did not induce hemolysis in blood, suggesting that these nanoparticles exhibit favorable intravenous safety profiles. Histopathological analysis via H&E staining revealed no significant organ toxicity following the final treatment (Fig. S7A). Furthermore, blood biochemical analyses indicated that all measured parameters remained within normal physiological ranges, supporting the biosafety and biocompatibility of CBP@HMnO_2_@HA (Fig. S7B-E). Collectively, these findings confirm that CBP@HMnO_2_@HA possesses a well-tolerated safety profile.

## 4. Conclusion

In conclusion, the ICD and cGAS-STING pathways have garnered significant attention in the field of tumor immunotherapy and are known to effectively enhance anti-tumor immune responses [31, 32]. Recent studies have demonstrated that chemotherapeutic agents such as CPT, cisplatin, carboplatin, and Mn^2+^ can induce immunogenic cell death (ICD) and activate the cGAS-STING signaling pathway [33, 34]. Therefore, this study developed a hollow mesoporous manganese dioxide (HMnO_2_)-based tumor-targeting nanomedicine (CBP@HMnO_2_@HA) designed to synergistically improve therapeutic outcomes in ovarian cancer by integrating chemotherapy (carboplatin) with chemodynamic therapy (Mn^2+^ -mediated Fenton-like reaction). The nanosystem employs HA for tumor-targeted delivery, releasing carboplatin within tumor cells to induce DNA damage and trigger ICD. Concurrently, Mn^2+^ generated during decomposition catalyzes the production of reactive oxygen species (•OH), thereby activating the cGAS-STING pathway, which further enhances ICD effects and initiates robust anti-tumor immune responses. The innovation of this strategy is threefold: (1) synergistic anti-tumor effects through combined chemotherapy and chemodynamic therapy; (2) dual activation of the STING pathway by carboplatin and Mn^2+^, leading to improved tumor immune microenvironment; and (3) HA surface modification that enhances targeting specificity and biocompatibility. This approach presents a promising solution to address the clinical challenges associated with high recurrence rates and poor survival outcomes in ovarian cancer. In vitro experiments demonstrated potent anti-tumor activity and effective induction of ICD. In vivo studies confirmed strong anti-tumor efficacy and activation of the cGAS-STING signaling pathway in an ovarian cancer mouse model. Nevertheless, further research is required to optimize drug delivery efficiency as well as the activation capacity of both ICD and cGAS-STING pathways. Our findings exemplify a nano-delivery platform that integrates carboplatin with HMnO_2_-mediated ROS generation strategies to activate ICD and cGAS-STING pathways both in vitro and in vivo, offering novel insights into enhancing chemoimmunotherapy and paving the way for future translational applications in anti-tumor therapy.

---

**Scheme 1.**
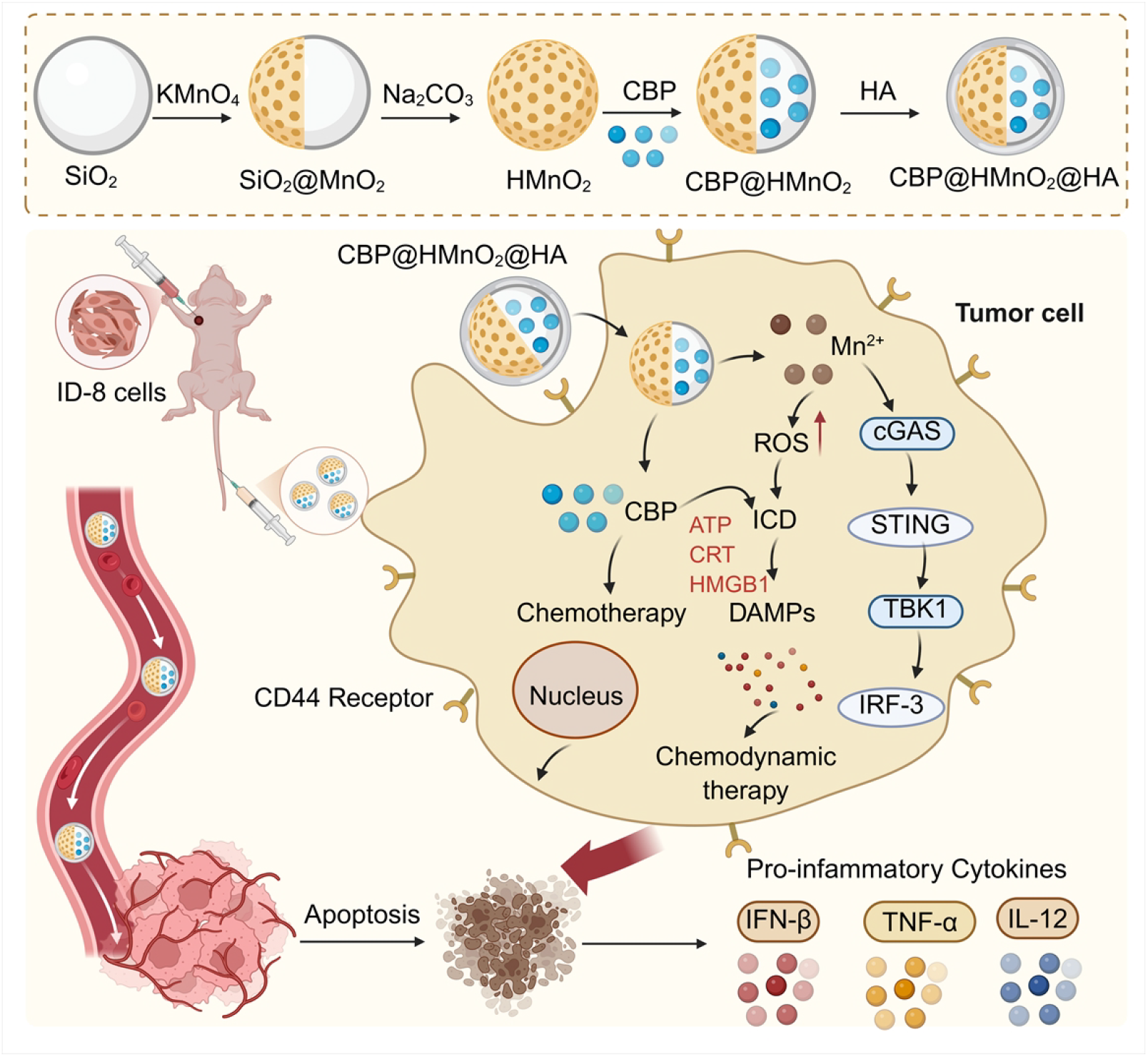
Schematic diagram of the preparation and biological functions of nanoparticles.

**Fig. S1.**
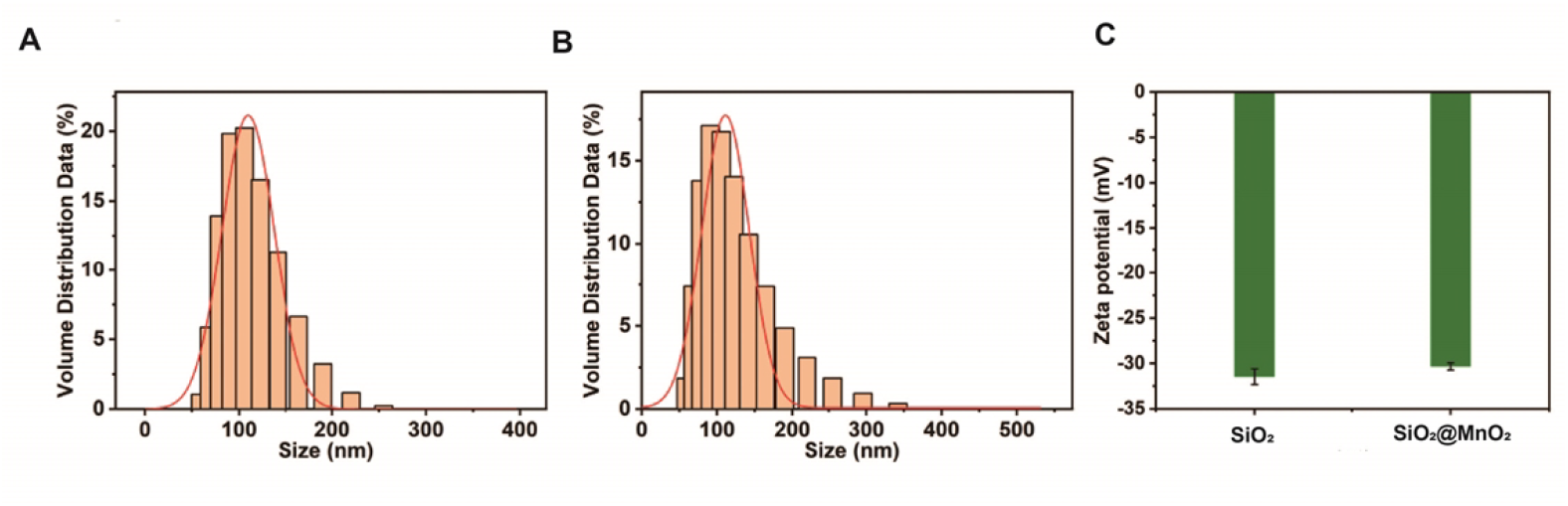
(A) The hydrated particle size of SiO_2_ nanoparticles. (B) Hydration particle size of SiO_2_@MnO_2_. (C) The Zeta potential of SiO_2_ and SiO_2_@MnO_2._

**Fig. S2.**
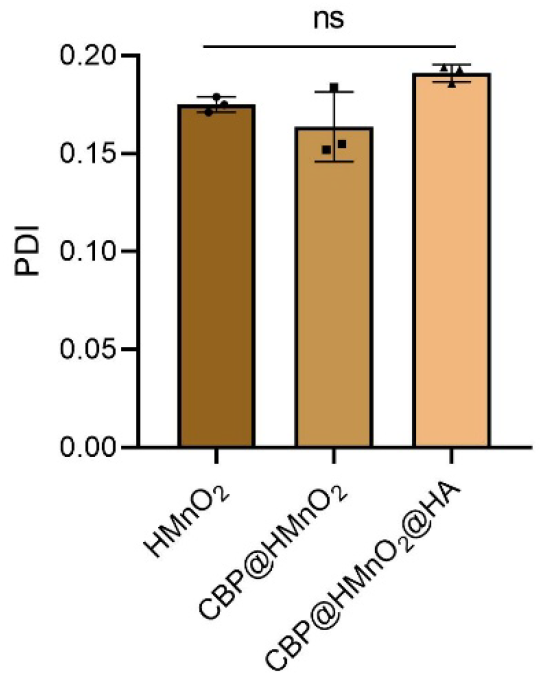
The PDI of HMnO_2_, CBP@HMnO_2_ and CBP@HMnO_2_@HA. Data are presented as mean ± SD, n=3. Statistical significances between every two groups were calculated via one-way ANOVA. *p < 0.05, **p < 0.01, ***p < 0.001, ****p < 0.0001.

**Table. S1.** The drug loading capacity and encapsulation efficiency of nanoparticles.

| DL (CBP@HMnO<sub>2</sub>@HA) | EE (CBP@HMnO<sub>2</sub>@HA) | DL (CBP@HMnO<sub>2</sub>) | EE (CBP@HMnO<sub>2</sub>) |
| --- | --- | --- | --- |
| 6.21 | 49.66 | 9.13 | 50.216 |
| 6.35 | 50.85 | 9.53 | 52.654 |
| 6.40 | 51.28 | 9.11 | 50.127 |

**Fig. S3.**
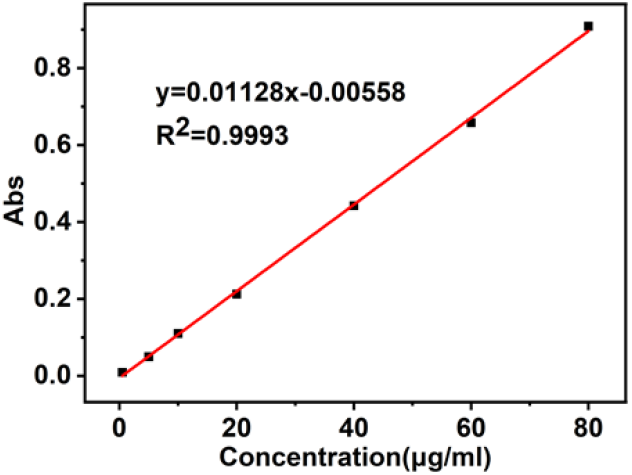
The standard curve of CBP (UV, 229 nm).

**Fig. S4.**
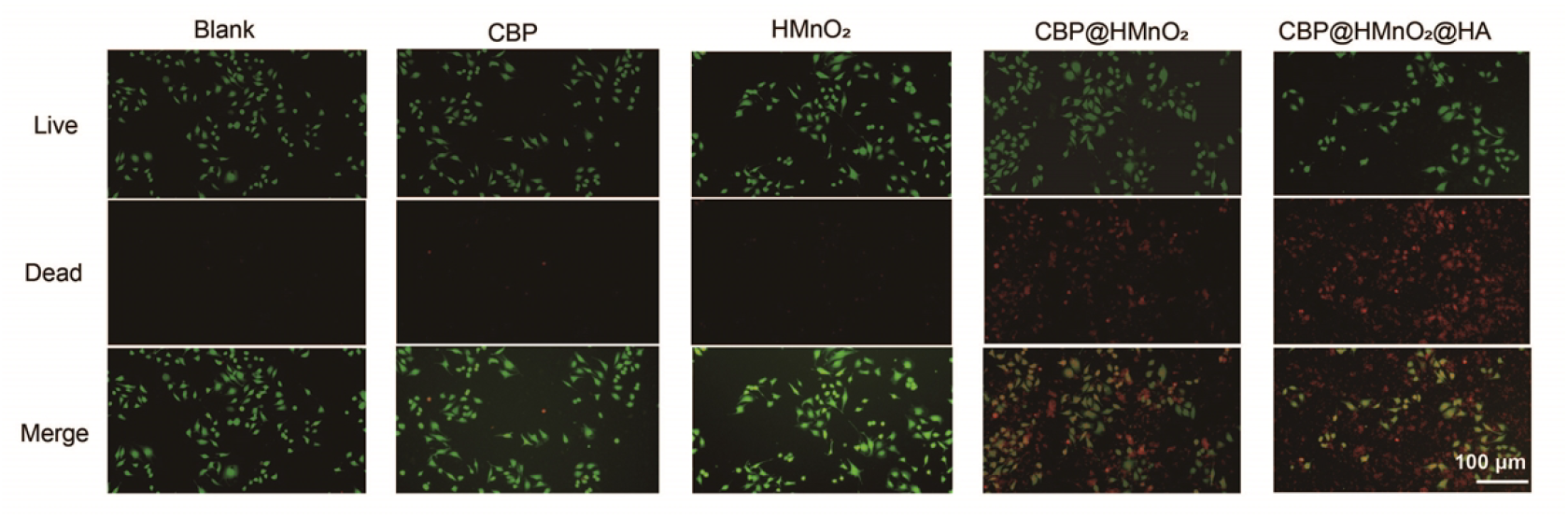
The live/dead staining assay of ID-8 cells treated with various drugs to evaluate cellular viability and cytotoxicity (scale bar represented 100 μm).

**Fig. S5.**
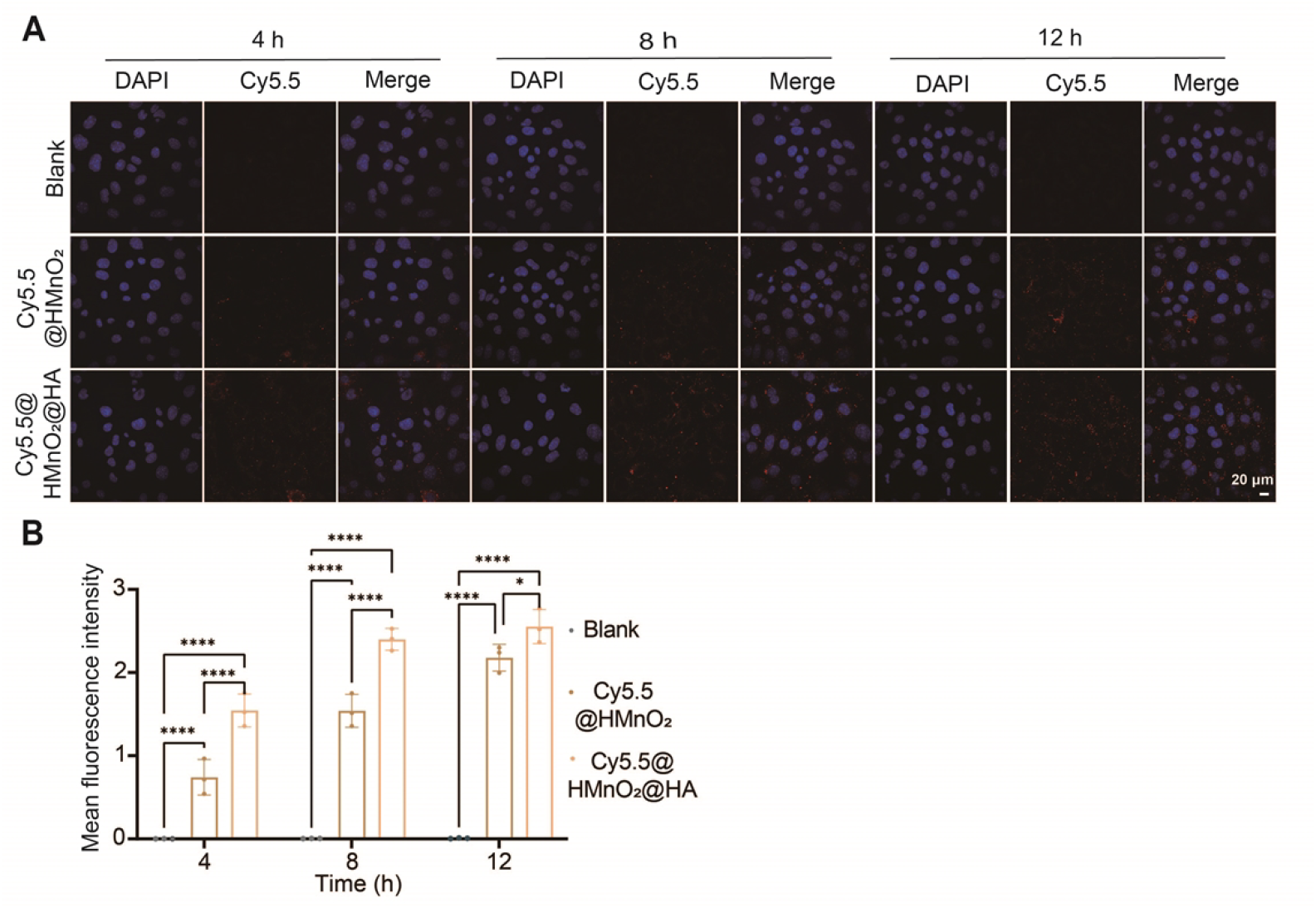
(A) Representative CLSM images of ID-8 cells treated with Cy5.5@NPs at 4 h, 8 h and 12 h, respectively. The Cell nucleus is strained by DAPI with blue fluorescence. The red fluorescence comes from Cy5.5 (scale bar represented 20 μm). (B) Quantitative Data Plot for Cellular Uptake of Nanoparticles at Various Time Points. Data are presented as mean ± SD, n=3. Statistical significances between every two groups were calculated via one-way ANOVA. *p < 0.05, **p < 0.01, ***p < 0.001, ****p < 0.0001.

**Fig. S6.**
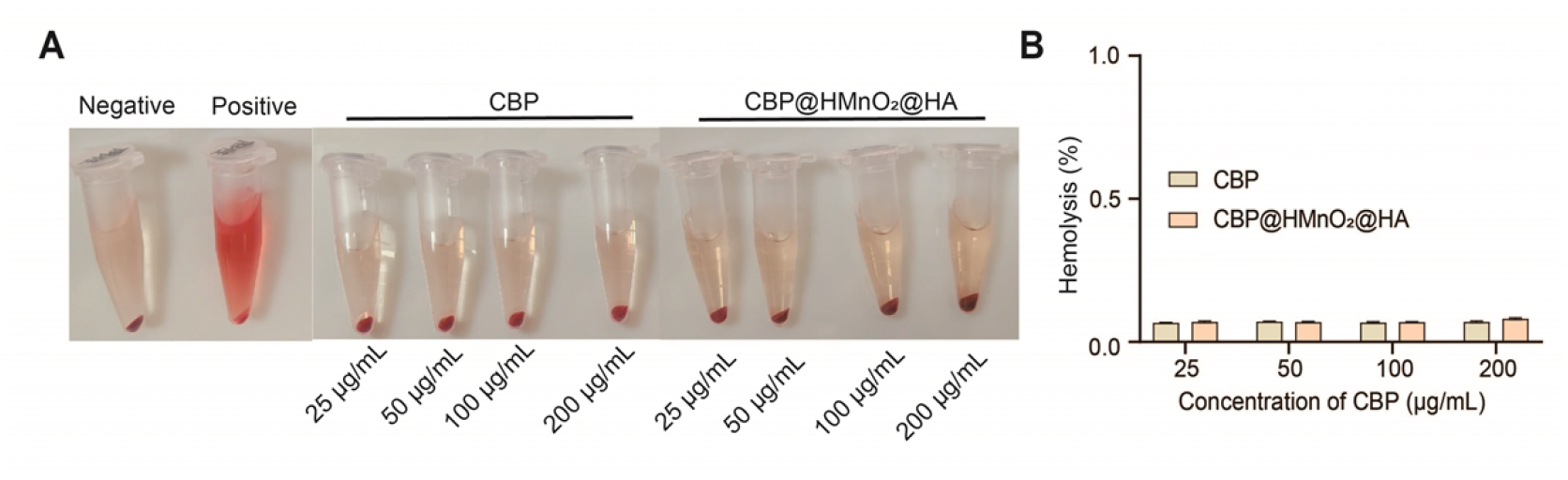
Apparent (A) and analytical (B) maps of hemolysis at different CBP and CBP@HMnO_2_@HA doses. Data are presented as mean ± SD, n=3. Statistical significances between every two groups were calculated via one-way ANOVA. *p < 0.05, **p < 0.01, ***p < 0.001, ****p < 0.0001.

**Fig. S7.**
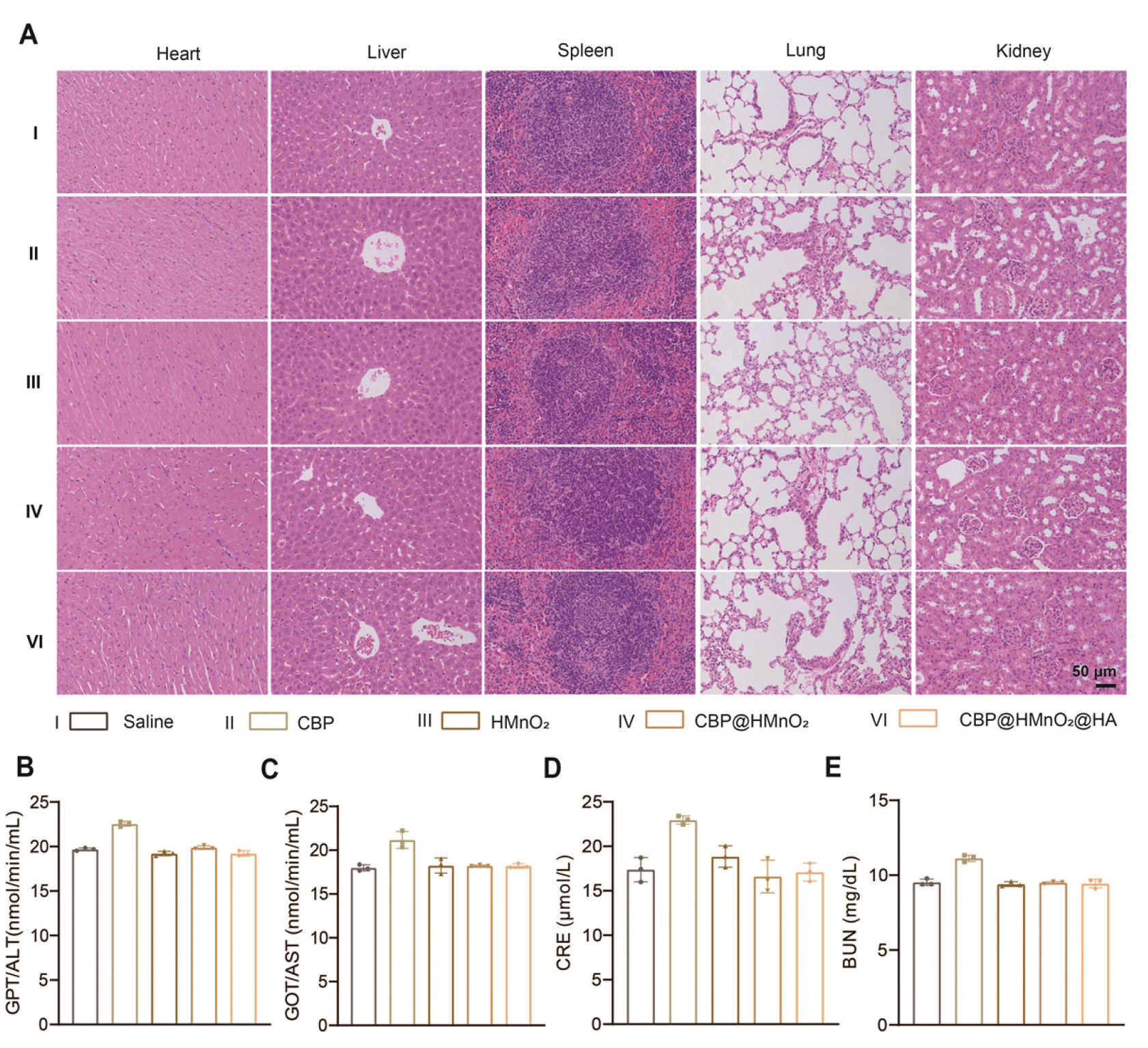
*In vivo* biosafety evaluation of NPs. (A) H&E staining of different groups of heart, liver, spleen, lung, and kidney (scale bars represented 50 μm). Serum ALT (B) and AST (C) levels at different groups. Serum levels of CRE (D) and BUN (E) at different groups. Data are presented as mean ± SD, n=3. Statistical significances between every two groups were calculated via one-way ANOVA. *p < 0.05, **p < 0.01, ***p < 0.001.

## Author contributions

Yong Wang: Writing – original draft, Methodology, Funding acquisition, Data curation, Conceptualization. Jie Pi: Project administration, Methodology, Writing – review & editing. Yuzi Zhao: Project administration, Software, Methodology. Qingzhen Xie: Supervision, Funding acquisition, Project administration.

## Appendix A. Supplementary data

Supplementary data to this article can be found online at X.

## Acknowledgements

The authors genuinely thank the management of Renmin Hospital of Wuhan University for their invaluable support and provision of the necessary resources to carry out this research work.

## Data availability

The authors declare that the data supporting the findings of this study are available within the paper and its Supplementary Information files. Should any raw data files be needed in another format they are available from the corresponding author upon reasonable request. Source data are provided with this paper.

